# Advancing Virus-Induced Gene Silencing in Sunflower: key factors of VIGS spreading and a novel simple protocol

**DOI:** 10.1101/2023.12.12.571232

**Authors:** Majd Mardini, Mikhail Kazancev, Elina Ivoilova, Victoria Utkina, Anastasia Vlasova, Yakov Demurin, Alexander Soloviev, Ilya Kirov

## Abstract

Virus-Induced Gene Silencing (VIGS) is a versatile tool in plant science, yet its application to non-model species like sunflower demands extensive optimization due to transformation challenges. In this study, we aimed to elucidate the factors that significantly affect the efficiency of Agrobacterium-VIGS in sunflowers. After reaffirming a number of approaches, we concluded that the seed vacuum technique followed by 6 h of co-cultivation produced the most efficient VIGS results. Genotype-dependency analysis revealed varying infection percentages (62-91%) and silencing symptom spreading in different sunflower genotypes. Additionally, we explored the mobility of tobacco rattle virus (TRV) and phenotypic silencing manifestation (photo-bleaching) across different tissues and regions of VIGS-infected sunflower plants. We showed the presence of TRV is not necessarily limited to tissues with observable silencing events. Finally, time-lapse observation demonstrated a more active spreading of the photo-bleached spots in young tissues compared to mature ones. This study not only offers a robust VIGS protocol for sunflowers but also provides valuable insights into genotype-dependent responses and the dynamic nature of silencing events, shedding light on TRV mobility across different plant tissues.

## 1. Introduction

Virus-Induced Gene Silencing (VIGS) has emerged as a powerful tool in many areas of crop research, including functional gene analysis [^1,2,3^], pathogen resistance studies [4, 5,^6^], abiotic stress tolerance [^7, 8, 9^], and general crop improvement [^10^]. The number of search projects dedicated to the VIGS technology is booming, highlighting its widespread adoption and recognition as a versatile tool for advancing crop science and biotechnology. An important factor for achieving high VIGS efficiency is the delivery method, which is commonly facilitated by Agrobacterium [^11^]. Unconventional delivery methods have also been established for model species, such as the use of cell-penetrating peptide carriers [^12^] and direct inoculation [^13^] in *Arabidopsis thaliana*, biolostic delivery [^14^], sap inoculation [^15^], carbon nanocarriers in *Nicotiana benthamiana* [^16^], and conjugated polymer nanoparticles in *N. tabacum* [^17^]. However, integrating VIGS systems could be challenging for non-model and recalcitrant species such as sunflowers.

One of the most challenging obstacles in VIGS studies is the adaptation of infection protocols to new species with high silencing efficiency. VIGS efficiency can be influenced by a multitude of factors. These factors encompass variables associated with the growth conditions of treated plants, such as photoperiod [^18^], humidity [^19^], and temperature [^20^]. Agrobacterium inoculation techniques also play a crucial role in VIGS efficiency [^21^]. An unconventional agrobacterium-mediated VIGS technique, which has been used in many challenging species, is vacuum infiltration at early developmental stages, such as seeds, germinated seeds, or sprouts. This approach was successful in wheat [^22^], Miscanthus [^23^], tomato [^24^], *Solanum rostratum* [^25^], *Solanum pseudocapsicum* [^26^], and *Lycium barbarum* [^27^]. However, the silencing efficiency of this technique may vary and is highly affected by different technical parameters, including bacterial concentration (OD_600_), vacuum and co-cultivation duration, and sprout size [^22, 24, 28^]. Another issue affecting VIGS efficiency is genotype dependency, which has been observed in soybean [^29^], cassava [^30^], citrus [^2^], and wheat [^31^].

Sunflower (*Helianthus annuus* L.) is a widely cultivated crop mainly used for the production of oil and confectionary products [^32, 33^]. Despite its agricultural significance, it has been traditionally considered a challenging species for transformation [^34^, ^35^, ^36^, ^37^]. In the field of VIGS, only a few studies on sunflowers have been conducted recently. In one study [^38^], authors used to validate the role of the Ha-ROXL gene in flower development. Two different infiltration techniques were used in this study: injection of the abaxial epidermis using a needleless syringe and wrapping a cut or scratched tissue with a piece of cotton soaked with infiltration suspension. In another study [^39^], authors successfully silenced the HaTubulin and OcQR1 genes using a seed soaking method, which required a pretreatment procedure of seeds surface sterilization and a post infection recovery on Murashige and Skoog medium for about 3 days. The moderate silencing efficiency and requirement of an in vitro culture step significantly limits the application of the VIGS technique in sunflower studies. Therefore, a robust and simple VIGS protocol needs to be developed for sunflowers.

Herein, we provide a simple seed-vacuum VIGS protocol for sunflowers (dx.doi.org/10.17504/protocols.io.261ged56dv47/v1). Apart from peeling the seed coats, our protocol requires no additional preparation of plant material. Notably, no *in vitro* recovery or surface sterilization steps were required. A simple seed-vacuum infiltration followed by 6 hours co-cultivation gave the most efficient VIGS results in terms of the infection percentage (up to 77%) and silencing efficiency of the targeted gene (normalize relative expression = 0.01). Another aspect we investigated in this study was the efficiency of VIGS in different genotypes, an issue that has not yet been addressed in sunflowers. Using our optimized VIGS protocol, we infected 6 different genotypes. Susceptibility to TRV VIGS infection and spread varied among the tested genotypes. The genotype ‘Smart SM-64B’ showed the highest number of infected plants (91%); however, the spreading of the silencing phenotype (the photo-bleached area) was the lowest in comparison to others. Finally, the mobility of TRV was investigated by RT-PCR in green and bleached tissues from all levels of infected plants. We found that the presence of TRV inside a VIGS-infected sunflower was not limited to tissues exhibiting a silencing phenotype, an observation similar to that reported for *Thalictrum dioicum* [^40^] and *N. benthamiana* [^41^]. More importantly, our results also showed that TRV was present at the highest level in the VIGS-infected sunflower (up to level 9), which indicates that our seed-vacuum protocol is able to facilitate extensive viral spreading throughout the infected plant.

## 2. Materials and Methods

### 2.1. Plant material and growing conditions

Seeds of sunflower lines ZS and Smart SM-64B were provided by All-Russia Research Institute of Oilseed Crops (Krasnodar, Russia) and “SMART” company (Krasnodar, Russia). From local seed providers (Ailita, Moscow; Dom Semyan, St. Petersburg), five different commercial sunflower cultivars were purchased: ‘Buzuluk,’ ‘Kubanski Semechki,’ ‘Lakomka, Shelkunshik,’ and ‘Oreshek.’ For the first two VIGS experiments (testing different infection methods, different growing phases, vacuum, and co-cultivation time) we used only sunflower seeds of line “ZS”. For the third VIGS experiment (testing VIGS efficiency on different genotypes) we used all five commercial cultivars in addition to line “Smart SM-64B”.

Infected sunflower plants were cultivated in a greenhouse using 7x7 cm plastic pots, with no gaps between pots within each treatment. The growth medium comprised a 3:1 ratio of peat to perlite. The greenhouse conditions were maintained at an average temperature of 22°C, with an 18-hour light and 6-hour dark photoperiod, and a relative humidity of approximately 45%. LED light system Just Grow X|Space (Grow Stuff, Moscow, Russia).

### 2.2. Constructing recombinant TRV vectors and agrobacterium transformation

The following TRV plasmids were used in our experiments: pYL192 (TRV1, Addgene #148968) and pYL156 (TRV2; Addgene #148969). A fragment of PDS (gene ID: 110913080. Assembly: HanXRQr2.0-SUNRISE) of *Helianthus annuus* L. was selected using pssRNAit software (https://www.zhaolab.org/pssRNAit/). Sunflower genomic DNA was used as a PCR template to amplify the VIGS insertion. The PCR reaction was conducted using a high-precision polymerase Tersus Plus PCR kit (Evrogen, Moscow, Russia) with the following primers: 5’-taattctagaATGGCATTTTTAGATGGCAGCCC-3’ and 5’-taatggatccTGGAGTAGCAAATACATAAGCATCCCC-3.’ The amplicons obtained were inserted into the XbaI and BamHI restriction sites of the TRV2 vector. Restriction reactions for plasmids and PCR amplicons were performed separately. Each reaction consisted of 1 μL of each endonuclease FastDigest (Thermo Scientific, USA), 2 μL of 10x FastDigest Buffer, 1 μg of DNA, and ddH2O to reach a total volume of 20 μL. Incubation was performed at 37°C for 2 hours followed by 5 minutes at 80°C to terminate the reaction. After purification using the Cleanup Standard kit (Evrogen, Moscow, Russia), the restriction products were ligated using T4 ligase (Evrogen, Moscow, Russia). The ligation reaction contained 100 units of T4 ligase, 2 μL 10× Overnight ligation Buffer, 50 ng plasmid + PCR amplicons (1:5 ratio respectively), and ddH2O to reach a total volume of 20 μL. The resulting recombinant plasmids were cloned into *the E. coli* strain dH5α. The obtained clones were grown on LB agar plates containing 50 μg/mL kanamycin.

All TRV constructs (pTRV1, pTRV2-empty, and pTRV2-HaPDS) were transformed into *Agrobacterium tumefaciens* (strain GV3101) using a standard electroporation procedure. Glycerol stocks of transformed *A. tumefaciens* were prepared and stored at -80°C for later use in VIGS infection experiments.

### 2.3. Agrobacterium cultures preparation and VIGS infection procedures

#### 2.3.1. Preparation the infiltration suspension

Frozen (−80C) 10% glycerol stocks of transformed agrobacterium were streaked on an LB-agar plate containing 10 μg/mL gentamicin, 50 μg/mL kanamycin, and 100 μg/mL rifampicin. The plates were incubated at 28 °C for 1.5 days. Two random single colonies were PCR-checked using the primers: for TRV1 R–AGACAACTTAATAACACATTGCGGACG, F – CTTTGACGTTGGAGTCCACGTTC; for TRV2-empty and TRV2-PDS(Ha) F–GTTCAGGCGGTTCTTGTGTGTC and R–TTGAACCTAAAACTTCAGA-CACGGATCTAC, which produced a PCR band of 234 bp for TRV2-empty and 417 bp for TRV2-PDS(Ha). Agrobacterium starting culture was obtained by inoculating PCR-validated colony in 5 mL LB medium with antibiotics (50 μg/mL kanamycin + 10 μg/mL gentamycin + 100 μg/mL rifampicin) and growing for 1.5 days in shaker at 28°C and 180 rpm. To prepare the infiltration culture, the starting culture was transferred to a larger flask with 45 mL liquid LB containing antibiotics in addition to MES (10 mM) and acetosyringone (200 μM). The infiltration culture was grown until an OD_600_ value of 1.5, which typically took about 5-6 hours. Later, agrobacterium was washed twice in sterile distilled water and resuspended in the infiltration buffer, which contained MES (10 mM), acetosyringone (200 μM), and MgCl2 (10mM). The obtained infiltration agrobacterium suspension was incubated in room temperature for 2 hours before performing VIGS infection. For VIGS infection, a 1:1 mix of TRV1:TRV2 agroba cterium suspensions was prepared.

#### 2.3.2. Syringe agroinfiltration

Sunflowers were grown in the greenhouse under an 18-hour light/6-hour dark cycle at 21°C until reaching the third true-leaf phase. The 1:1 TRV1:TRV2 agrobacterium mixture was prepared as mentioned above and then injected into the abaxial epidermis of each leaf using a needleless medical syringe (each plant requires ≈0.5 mL of the 1:1 mixture). The infiltrated plants were covered with plastic wrap and incubated in darkness at room temperature for 2 days. Subsequently, the plastic wrap was removed and the plants were returned to the greenhouse.

#### 2.3.3. Rubbing

The rubbing technique was also employed on sunflowers during the third true-leaf phase. 2 g of autoclaved silica sand was introduced into a beaker containing 10 mL of the 1:1 TRV1:TRV2 mixture (sufficient for infecting 10-12 plants using this method). After brief stirring, the sand-agrobacterium combination was gently rubbed onto the surface of the sunflower leaves. The plants were subsequently covered with plastic wrap and placed in darkness at room temperature for 2 days before being returned to the greenhouse.

#### 2.3.4. Seed-vacuum infiltration

The outer coat of the sunflower seeds was removed, and the seeds were subsequently soaked in tap water for 0.5-1 hour to facilitate the removal of the inner coat. Afterward, 12-15 pealed seeds were placed into a Petri dish along with 20 mL of the 1:1 agrobacterium mixture and 4 g of autoclaved silica sand. Using the tips of an index finger, the seeds were gently scrubbed against the surface of the Petri dish in a circular motion for 30 seconds. Care was taken to change the gloves between treatments. The Petri dishes were then placed in a vacuum chamber. Negative pressure was applied for 3 minutes, followed by a 3-minute pause, and another 3 minutes of negative pressure. The Petri dishes were sealed with Parafilm and incubated in a shaker at 28°C and 50 rpm for 2, 6, or 18 hours. After incubation, the seeds were planted in pots containing humid soil and covered with a plastic wrap. Pots were kept in the dark at room temperature for 2 days, after which the plastic covers were removed, and the pots were relocated to the greenhouse. VIGS symptoms began to appear within 11-13 days. The same procedure described here was performed on 2-day-old sprouts. An overview of the VIGS procedure is presented in Figure 2. Extended technical details and other bench notes are published in a step-by-step protocol: dx.doi.org/10.17504/protocols.io.261ged56dv47/v1.

### 2.4. RT-PCR analysis

Reverse transcription-PCR analysis (RT-PCR) was conducted to assess TRV infection and its systematic spread. We extracted total RNA from a 50-100 mg leaf tissue sample using the ExtractRNA kit (Evrogen, Moscow, Russia), based on TRIzol reagent. DNase treatment was performed to remove any remaining DNA from the samples. cDNA synthesis and PCR amplification took place in the same reaction mix (one-step RT-PCR). The procedure was performed using the BioMaster RT-PCR – Premium (2×) kit (Biolabmix, Novosibirsk, Russia). The primers were designed to flank PDS insertion in the TRV2 construct: TRV2-HaPDS F – GTTCAGGCGGTTCTTGTGTGTC and R – TTGAAC-CTAAAACTTCAGACACGGATCTAC. 1 μl of DNase-treated RNA (≈100 ng) was used as a template. The reaction conditions were as follows: cDNA synthesis at 45°C for 30 minutes, followed by PCR with an initial denaturation step at 93°C for 5 minutes, and then 35 PCR cycles consisting of 15 seconds denaturation at 93°C, 20 seconds annealing at 60°C, 20 seconds elongation at 68°C, and 5 minutes final elongation at 68°C. RT-PCR results were visualized in 2% agarose gel.

### 2.5. Expression analysis of the VIGS target gene

Quantitative PCR (qPCR) was used to evaluate changes in expression levels of PHYTOENE DESATURASE (PDS) upon VIGS infection. Total RNA extraction was performed as shown in the previous section. For cDNA synthesis, DNase-treated RNA was primed with oligo(dT)10 primer. The reaction was performed using the MMLV RT kit (Evrogen, Moscow, Russia) according to the manufacturer’s instructions. qPCR was performed using BioMaster HS-qPCR SYBR Blue(2×) kit (Biolabmix, Novosibirsk, Russia). To evaluate changes in PDS expression levels, primers were designed specifically to amplify a region of the gene that did not include the VIGS insertion: Ha-PDS-qPCR-F – ATCCGAATCGGTCAATGTTGG and R – TGCTCCCATCTTCTGCAATTT. Additionally, qPCR primers for Actin of *Helianthus annuus* were designed to be used as reference. The amplification reaction was conducted via Bio-Rad Real-time CFX96 Touch cycler. The qPCR conditions were: initial denaturation step at 95°C for 5 minutes; 40 cycles consisting of 15 seconds denaturation at 95°C and 15 seconds annealing + elongation at 58°C; plate read was conducted at the end of each cycle. Data analysis and gene expression profiling were performed using BioRad CFX Manager™ Software. The ΔΔCq mode was used for normalization.

### 2.6. Phenotyping of photo-bleaching symptoms

A total of 77 sunflower plants were evaluated for key traits, including the number of levels, the number of symptomatic leaves (SL), the number of all leaves (AL), and the position of the uppermost symptomatic leaf. The collected data were used to calculate various parameters for each treatment, such as the percentage of symptomatic plants to all plants, the number of plants with symptomatic apical leaves, and the average ratio of symptomatic leaves to all leaves (SL/NL). To maintain consistency in the dataset, specific guidelines were established for phenotyping: The determination of the number of levels in a given plant was based on the growing symmetry, since sunflowers might sometimes exhibit double or triple symmetry. The apical leaf was identified as the highest recognizable leaf with a minimum length of 0.5 cm. Symptomatic leaves were defined as those displaying at least one bleached spot, regardless of the spot’s diameter.

For the VIGS experiment where we tested different genotypes susceptibility we used ImageJ software [^42^] to measure the percentage of photo-bleached area. The color threshold was set to register only intensely white pixels. This means that white-green or white-yellow areas were not included in the calculations of the photo-bleaching percentage.

## 3. Results

### 3.1. The effect of different infection methods and plant growth phases on VIGS efficiency of sunflower PDS gene

To determine the optimal conditions for VIGS we tested the influence of plant growth phase (plants at the 3rd true leaf stage, 2-day-old sprouts, or seeds) and the technique of Agrobacterium infection (syringe, rubbing, or vacuum) on the tobacco rattle virus (TRV) mediated VIGS efficiency. We used the phytoene desaturase (PDS) gene as a control for VIGS efficiency because of the clear photo-bleaching phenotype resulting from PDS silencing. The selected fragment of the sunflower PDS gene was cloned into the TRV2 vector and used for further analysis. The following variants were used in this experiment: (1) syringe infiltration during 3rd-true-leaf phase; (2) rubbing during 3rd-true-leaf phase; (3) vacuum-infiltration of 2-day-old sprouts followed by 2-hr co-cultivation; (4) vacuum-infiltration of seeds followed by 2-hr co-cultivation. Each approach was performed using three VIGS variants: TRV1 + TRV2-empty, TRV1 + TRV2-PDS(Ha), and infiltration buffer as a negative control.

Syringe infiltration, rubbing, and vacuum infiltration of sprouts showed very little to no silencing events. Only 2 out of the 12 syringe-infiltrated plants (approach 1) showed very mild photo-bleaching symptoms. Rubbing of leaves and vacuum infiltration of sprouts (approaches 2 and 3) showed no photobleaching symptoms. Out of the 15 vacuum-infiltrated seeds (approach 4), 13 developed into adult plants, and 8 of them (61%) displayed photo-bleaching symptoms. VIGS symptoms began to appear between 11-13 dpi (days post infection). Plants treated with TRV without gene inserts (empty TRV) had no photo-bleaching symptoms, but possessed characteristic TRV symptoms, that is, leaf curling and yellowing (Figure 1A).

**Figure 1.**
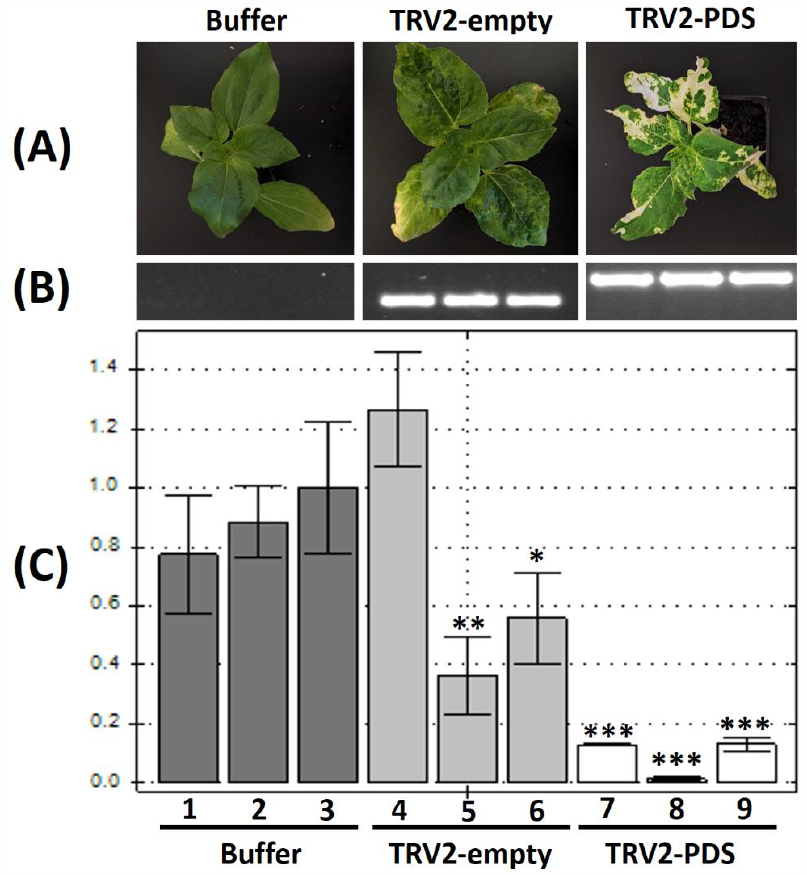
VIGS of the sunflower PDS gene performed by the seed vacuum-infiltration approach: (**A**) Representative sunflower plants from each VIGS variant: Buffer, TRV2-empty, TRV2-PDS(Ha); (**B**) RT-PCR results with primers designed on the regions of TRV2 RNA flanking the insertion site. From left to right: Buffer treatment as negative control – the total absence of PCR bands indicates no cross-contamination during the experiment; TRV2-empty – PCR band with the expected length (234bp) indicates the presence of the corresponding TRV RNA; TRV2-PDS(Ha) – PCR band with the expected length (417bp) indicates the presence of the corresponding TRV; (**C**) The qPCR results of PDS gene expression for the same samples as in (B) (ordered respectively). The Y-axis shows the value of relative normalized expression. The normalization mode used – ΔΔCq. The control was set to sample number 3 (buffer sample), indicating the 1.0 value of normalized relative expression. Each bar shows the mean±SD for the relative normalized expression of the 3 technical replicates. *T*-Test was performed between all biological replicates and the control sample. Samples with significant difference are marked with asterisks (*p < 0.05, **p < 0.01, ***p < 0.001).

To further validate the VIGS results, we conducted RT-PCR and qPCR analyses for TRV RNA detection and PDS gene expression estimation, respectively. Plant materials were collected only from the seed vacuum infiltration group (approach 4). Total RNA was extracted from three plants (three biological replicates) for each VIGS treatment (TRV2-empty, TRV2-PDS, and Buffer). RT-PCR confirmed the viral presence in all the symptomatic plants we examined, as evidenced by distinct PCR bands observed in TRV2-PDS and TRV2-empty variants (417 bp and 234 bp, respectively (Figure 1B)). As expected, no PCR bands were detected in the control (buffer infiltration) plants (Figure 1B). The same RNA samples were used for qPCR analysis of PDS expression. Three technical replicates were performed for each sample. Compared to the control, the relative normalized expression of PDS was significantly lower in the TRV2-PDS treated plants compared than in the TRV2-empty and control (buffer infiltration) variants (Figure 1C). The PDS expression level in the three plants treated with the TRV2-empty construct was also lower than that in the control. Interestingly, one of the biological replicates of the TRV2-empty treatment showed an increase in PDS expression with a relative normalized expression value of 1.26 (SD = 0.192) (Figure 1C).

Thus, the results show that the method of infection of sunflower plants by Agrobacterium and the plant growth stage significantly influence the VIGS efficiency. We found that vacuum infiltration of dry seeds followed by a period of co-cultivation resulted in the highest number of sunflower plants with VIGS symptoms.

### 3.2. The effect of Agrobacterium co-cultivation time and vacuum application on VIGS efficiency

We next tested the effect of the presence/absence of vacuum and the time of co-cultivation of sunflower seeds and Agrobacterium suspension. The experiment involved groups with and without the application of vacuum. In each group, there were three variants of post-vacuum Agrobacterium-seed co-cultivation time: 2, 6, or 18 h. Photobleaching symptoms were used to control the VIGS efficiency. We found that 75%, 76% and 71% of plants showed PDS silencing symptoms in the variants with 2, 6 and 18 hours of post-vacuum co-cultivation, respectively. In the group where no vacuum was used, 69%, 36%, and 20% of plants showed photo-bleaching symptoms in variants with 2, 6 and 18 hours of co-cultivation, respectively. Thus, the obtained results showed that vacuum application can improve the efficiency of VIGS for sunflowers (Table 1).

**Table 1.**
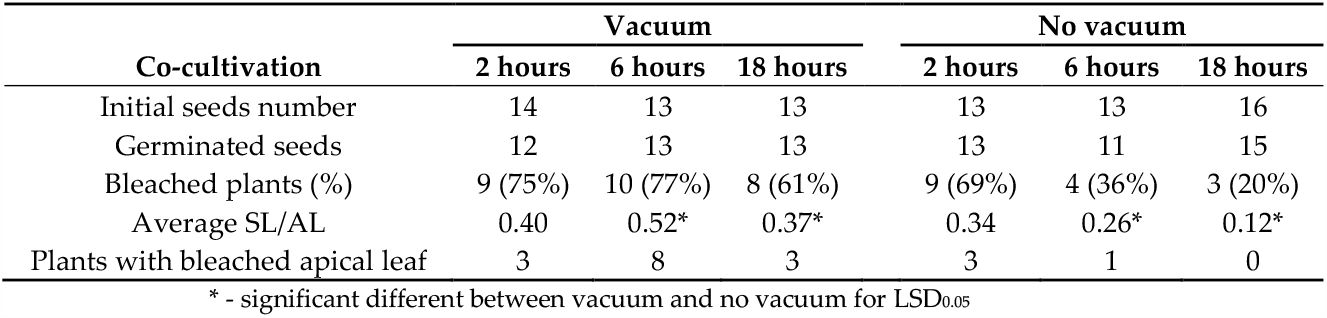
Effect of vacuum and co-cultivation time on the efficiency of Agrobacterium-VIGS infection of sunflower seeds.

Next, we assessed the uniformity of the VIGS symptoms at the whole-plant level. For this, we collected data on the number of Symptomatic Leaves (leaves exhibiting photo-bleaching symptoms) relative to the total number of leaves (SL/AL ratio). The SL/AL ratio ranged from 0 (no photobleaching) to 1 (all leaves showed photobleaching symptoms). A difference in the SL/AL ratios between the vacuum and no-vacuum treatments was evident (Table 1). The average SL/AL at 6-hour co-cultivation was 0.52 for the vacuum group and 0.26 for no vacuum (LSD_0.05_ = 0.20). Similarly, the average SL/AL ratio at 18-hour co-cultivation was 0.37 and 0.12, respectively (LSD_0.05_ = 0.17).

Another parameter we considered for the assessment of VIGS intensity in relation to different methodological factors was the manifestation of photobleaching symptoms on apical leaves. This parameter holds particular significance for VIGS studies because it might provide an insight into TRV symptom stability and TRV transition towards the growing point. In all treatments, except for the vacuum + 6 hours co-cultivation, few plants showed photo-bleaching symptoms on the apical leaf. However, in the vacuum + 6 hours co-cultivation group, 8 out 10 symptomatic plants showed photo-bleaching on the apical leaves (Table 1 and supplementary Table S1).

Altogether, the obtained results showed that the vacuum agroinfiltration of dry sunflower seed followed by 6 hours of co-cultivation with Agrobacterium suspension proved to be the most efficient method for ensuring VIGS in sunflower. A summary of our seed-vacuum VIGS protocol is shown in Figure 2. The simplicity and minimal preparation requirements outlined in our protocol make it accessible for future VIGS experiments on sunflowers. Notably, the methodology we developed yielded consistent results when performed by different people in our laboratory, even under semi-sterile conditions. We firmly believe that when coupled with well-designed VIGS constructs, our method should reliably produce the expected gene-silencing outcomes. The detailed protocol is available at dx.doi.org/10.17504/protocols.io.261ged56dv47/v1.

**Figure 2.**
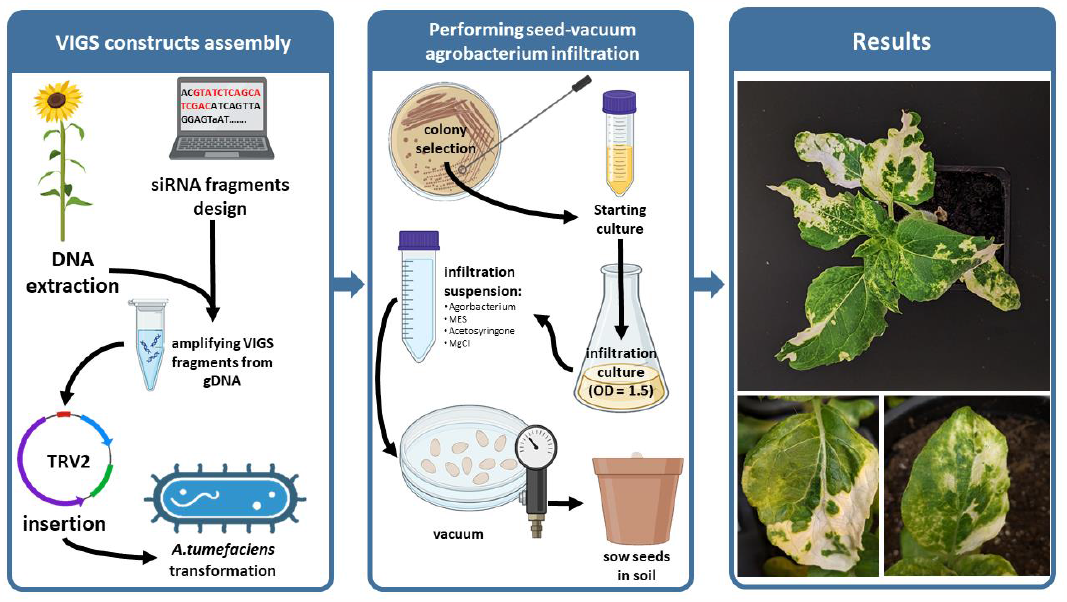
Overview of the seed-vacuum VIGS protocol.

### 3.3. Genotype dependency of VIGS in sunflower

To assess whether the optimized VIGS protocol is sufficient for different sunflower genotypes, we tested it on five sunflower cultivars and one breeding line. We evaluated the efficiency of the protocol by calculating the number of infected plants exhibiting photobleaching symptoms and the extent of the photo-bleached areas of leaves throughout the plant. Fourteen seeds from five commercial cultivars and one breeding line (see the materials section) were used for this experiment. Twenty-two days after agroinfiltration using the developed protocol, phenotyping data were gathered to calculate the following parameters: 1) percentage of plants with photo-bleaching symptoms (regardless of the intensity of the symptoms), 2) the SL/AL ratio, and 3) the percentage of bleached leaf area.

Differences in photobleaching intensity between the genotypes were noticeable (Figure 3). “Lakomka” and “Oreshek” exhibited pronounced symptoms with extensive spread of bleached spots across several individuals. Three other cultivars, ‘Shelkunshik,’ ‘Buzuluk’ and ‘Kubanski Semechki’ showed lower VIGS symptoms. Interestingly, the breeding line ‘Smart SM-64B’ showed mild spreading of photo-bleached spots (Supplementary Figure S1). However, when calculating the percentage of infected plants (plants with photo-bleaching symptoms in at least one leaf), genotype ‘Smart SM-64B’ exhibited the highest percentage, with 91% of plants infected. The percentages of infected plants in other genotypes were as follows: ‘Shelkunshik’ – 79%; ‘Buzuluk’ – 77%; ‘Kubanski Semechki’ – 67%; ‘Lakomka’ – 67%; ‘Oreshek’ – 62%. A single plant from each group, chosen based on the highest SL/Al ratio, was selected to calculate the percentage of leaf area affected by bleaching (Figure 3).

**Figure 3.**
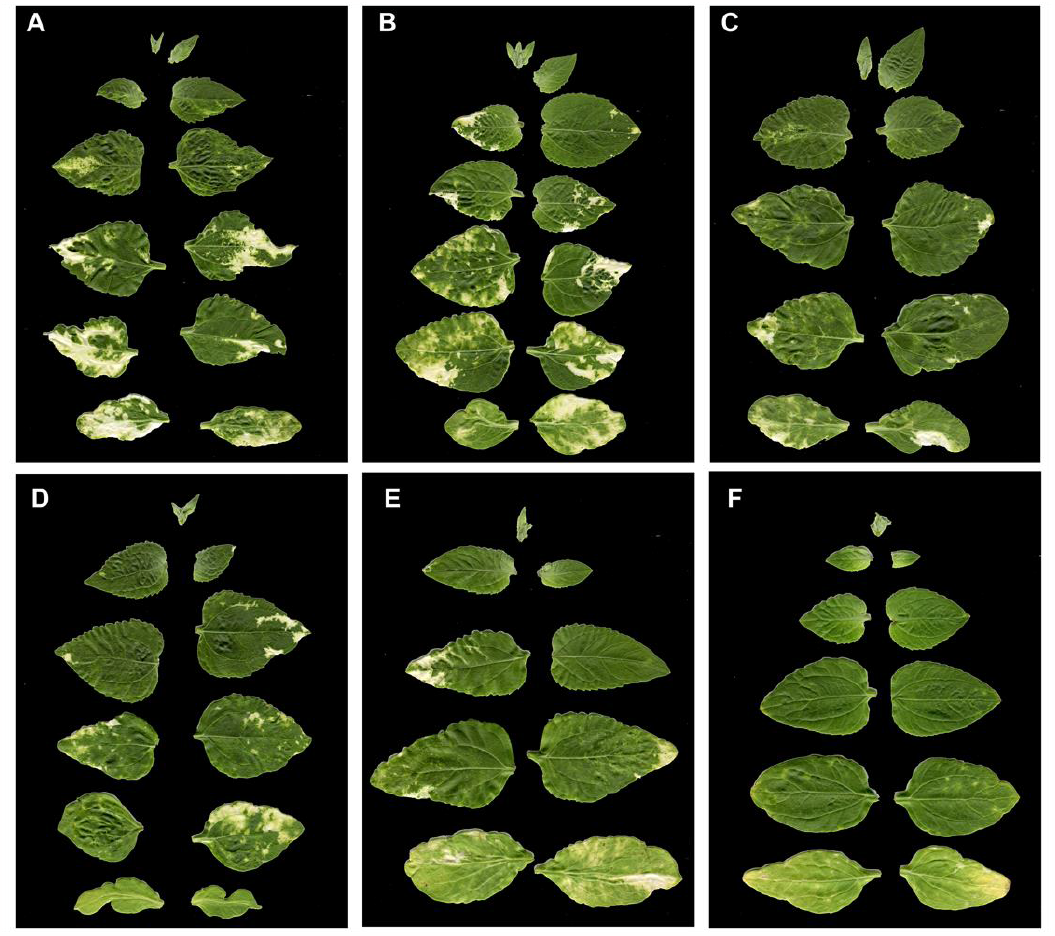
Scans of the leaf areas of plants with the highest SL/AL values from each genotype group. These images were individually used to calculate the percentage of bleached leaf area (% of bright white area of the whole leaves surface): (**A**) “Oreshek” – 16%; (**B**) “Lakomka” – 12%; (**C**) “Shelkunshik” – 5%; (**D**) “Buzuluk” – 4%; (**E**) “Kubanski Semechki” – 4 %; (**F**) “Smart SM-64B” – 3 %.

Thus, the optimized VIGS protocol showed good efficiency (62-91%) in terms of the number of infected plants for different sunflower genotypes. However, the rate of virus spread and the extent of VIGS symptoms vary significantly between genotypes.

### 3.4. TRV VIGS transmission from bottom to upper leaves

Viral movement throughout the plant system is an important factor to consider in VIGS studies. Elucidating viral dynamics plays a pivotal role in predicting which plant segments are likely to manifest silencing effects. Sunflower plants infected with the TRV2-PDS construct using the vacuum-seed method were transplanted into a larger pot to allow extensive growth and formation of new levels. At the age of 1 month, green and photo-bleached samples were collected from levels 2 to 7 for RNA extraction and RT-PCR analysis. Only green samples were collected from levels 8 and 9, as the phenotypic symptoms of PDS silencing were completely absent. We were unable to collect any samples from level 1 because the leaves were desiccated by that age.

According to the RT-PCR results, TRV was detected in the lower (level 2-3) as well as in top levels (level 8-9). We were unable to detect TRV at levels 6 and 7 (Figure 4b) for unknown reasons. However, these results were expected because of the weak PDS silencing phenotype on the leaves at these levels (Supplementary figure S2). We also separately analyzed the presence of TRV in the photo-bleached patches and the green parts of the leaves (Figure 4B). TRV was detected in both groups of samples, indicating that the presence of the virus does not always lead to VIGS symptoms. Interestingly, TRV was found at levels 8 and 9 despite the lack of any observable photobleaching symptoms, suggesting that a longer time is needed for VIGS symptom appearance.

**Figure 4.**
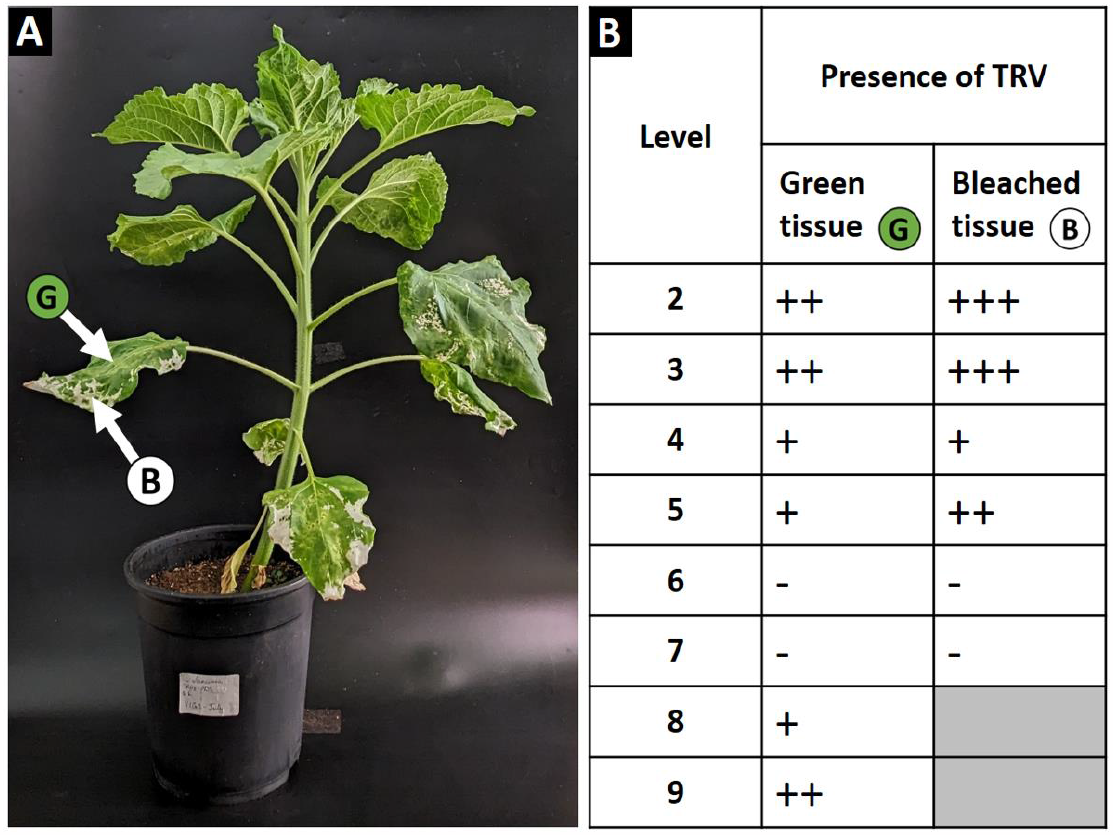
Tracking the presence of TRV in green and bleached tissues across VIGS-infected sunflower levels. (**A**) Sunflower plant one month after VIGS-PDS infection using the vacuum-seed method and 6 hours of co-cultivation (pot height: 20 cm). (**B**) RT-PCR results. RNA was extracted from the green and bleached tissues at different levels (starting from level 2) of the sunflower, as shown in A. The number of (+) signs represents the intensity of the RT-PCR bands, providing a semi-quantitative analysis of the viral load in a given sample. The (−) sign represents no RT-PCR; hence, complete viral absence. Electrophoresis and images of each plant material sample are provided in Supplementary figure S2.The results of these experiments showed efficient virus particle production after plant infection using our protocol, although VIGS symptoms did not always appear.

Furthermore, we conducted time-lapse observations to track the spread of photo-bleached spots in VIGS-infected sunflower plants. We captured an image of an infected plant every 30 min over a 4-day span. As demonstrated in Video S1, the silencing phenotype spreads in the actively growing tissues of young developing leaves. The photo-bleached spot exhibited an extension in size as the leaf grew. In mature, fully developed leaves of the same plant, the photo-bleached spot remains in a fixed state, with no observable changes in its shape or extension. This observation suggests a correlation between leaf developmental stage and the efficacy of gene silencing. The extension of the photo-bleached spot with leaf growth indicates a dynamic and ongoing silencing process, possibly related to the high level of permeability to viral particles in young tissues. Conversely, the fixed state of the photo-bleached spot in fully developed leaves might imply potential resistance to viral mobility in mature tissues.

Thus, our study demonstrates the successful implementation of the VIGS protocol for sunflowers, facilitating TRV transmission from the lower to the apical leaves. Additionally, this research presents a time-lapse observation capturing the spreading of the silencing phenotype across various tissues in VIGS-infected sunflowers. These observations provide novel insights into the mobility of TRV and the dissemination of the silencing phenotype within VIGS plants. Such examinations contribute to a better understanding of the dynamics involved in VIGS processes, offering valuable information for future studies in the field.

## 4. Discussion

Although subtle agrobacterium-mediated VIGS protocols have been well established for numerous species, a noticeable gap in technical details has become evident when applying this methodology to sunflowers. To address this gap, we initially sought to reproduce the VIGS protocols described in existing literature. However, we encountered several setbacks that may have been influenced by the sunflower genotypes used. Consequently, a series of optimization experiments was conducted to assess the influence of several key factors on VIGS efficiency: growth phase of the infected plant, infection technique, and genotype dependency. Each of these factors was tested for its effect on silencing efficiency, either alone or in combination with other factors. The optimization experiments resulted in a simple and rapid VIGS protocol that provided sufficient results in terms of the number of infected sunflower plants exhibiting VIGS symptoms, viral transmission to the apical leaves, and the level of target gene (PDS) silencing.

Regardless of other existing delivery approaches, such as direct infection with viral particles, *Agrobacterium* remains the method of choice in VIGS experiments [^43^]. Nonetheless, despite its apparent simplicity, the success or failure of agrobacterium-mediated VIGS strongly depends on the method employed for plant infections [^44^]. Surprisingly, application of the syringe-infiltration method and vacuum infiltration of germinated seeds did not lead to sufficient VIGS results. Notably, the most commonly used agroinfiltration method for VIGS is the syringe infiltration of leaves. This methods has been applied in VIGS studies on a broad range of plant species: *Brassica oleracea* [^3^], *Gerbera hybrid* [^45^], *Solanum lycopersicum* [^46^], lettuce [^47^], *Oryza sativa* [^48^], *Triticum aestivum* [^31^], *Eschscholzia californica* [^49^], *Capsicum annuum* [^50^], *Gossypium hirsutum* [^51^]. In sunflower [^38^], using syringe infiltration, the authors observed the photo-bleaching phenotype in expanded leaves and leaflets near the inflorescence. However, based on our experience, this method does not yield similar results. Using the syringe-infiltration technique, we only achieved mild photo-bleaching spots with no systematic spreading towards new leaves. In another sunflower VIGS study [^39^], Jiang *et al*. employed a seed soaking technique. This technique involved immersing sunflower seeds in the infiltration suspension for 6 hours, followed by 2 days of culturing in liquid MS medium. Afterward, the seeds were transferred to solid MS medium for 36 hours before being planted in soil. According to the authors, this technique resulted in 91% infection efficiency, as evidenced by the characteristic symptoms of TRV. Repeating this methodology in our laboratory resulted in significant difficulties during the MS cultivation stage. Despite the rigorous seed sterilization methods we employed, we consistently faced issues with contamination of MS media.

To the best of our knowledge, apart from the two studies mentioned above [^38, 39^], there are no VIGS protocols for sunflowers published in the current literature. Owing to our unsuccessful attempts to repeat the available methods, we embarked on a comprehensive series of experiments aimed at investigating the methodological factors that affect VIGS in sunflowers. As a result, we concluded that seed-vacuum infiltration followed by 6 hours of co-cultivation revealed to be the simplest and most effective approach. The seed-vacuum technique has already demonstrated its effectiveness in species that usually exhibit poor susceptibility to classical methods of *Agrobacterium* infiltration. In a study on maize and wheat [^22^], the authors showed a positive correlation between seed vacuum, cocultivation times, and silencing efficiency. In our study, we observed the same relationship: the overall percentage of symptomatic plants was higher in the vacuum treatments than in the no-vacuum treatments (Table 1). It was also shown previously that the vacuum infiltration technique was successful when applied to germinated seeds of wheat and tomato [^22, 24^]. However, the vacuum infiltration of the germinated seeds of sunflower (2-day-old sprouts) did not lead to any plants with photo-bleaching symptoms.

Another important aspect that we also studied in this work was the genotype response to VIGS in sunflowers. We showed that the choice of sunflower genotype could play a critical role in the efficiency and intensity of VIGS. While some sunflower cultivars might show a high percentage of plants with VIGS symptoms, the intensity of the symptomatic spreading of photobleaching can be very low (Figure 3). The same genotype dependency of VIGS was observed in several previous studies [^2, 29, 45^]. These observations highlight the significance of selecting appropriate plant materials for future VIGS research projects. When adapting VIGS systems for new, more complex targets, it must be more reasonable to initially work with susceptible genotypes in order to be able to evaluate the outcomes more effectively.

The final aspect related to VIGS that we investigated in our study was the mobility of TRV in various regions and tissues of infected plants. TRV belongs to the Tobravirus genus [^52^]. Viruses in this classification have two genomes, each consisting of a positive-sense single-stranded RNA (+ssRNA) [^53, 54^]. The bipartite nature of the TRV adds a layer of complexity while studying its behavior and movement inside the host. TRV has a unique capacity to enter the shoot apical meristem of plants [^55^]. This information is particularly important because it might provide insight into potential genetic changes in germline cells [^56^]. Our results showed that the presence of TRV in a VIGS-infected plant is not limited to the tissues where silencing events (photo-bleaching, in our case) were observed (Figure 4). We also observed a pattern between the duration of co-cultivation after vacuum treatment and the occurrence of silencing symptoms on the highest leaf in an infected plant (Table 1).

The observation we made regarding spreading of silencing phenotype could support various new insights for future VIGS studies. As we showed in Video S1 silencing spreads more actively in young leaves compared to mature ones. This phenomenon must be mainly linked to the inherent movement behavior of TRV within the host. It is widely acknowledged that the primary trafficking systems for viruses in plants involve plasmo-desmata [^57, 58^]. Heinlein [^59^] reported, that the size exclusion limit (SEL) in plasmodesmata is highly affected by the plant development stage and maturity. The structure of plasmo-desmata changes as the tissue grows from simple to more complex or “branched”, thus decreasing the SEL in mature leaves, which limits viral mobility and eventually the silencing activity.

## 5. Conclusion

In this study, we developed a simple and efficient protocol for VIGS in sunflowers. The main advantages of our protocol: (1) no pretreatment of the plant materials is required, such as sterilization of the surface of seeds; (2) no recovery step on *in vitro* medium is needed; (3) high efficiency of plant infection for different genotypes; (4) ensuring TRV transmission from lower to apical leaves. The new protocol for VIGS will be useful for functional genomics studies in sunflowers and other recalcitrant plants.

## Supporting information

Figure S1: VIGS infection in various sunflower genotypes. The red asterisk identifies plants used to calculate the percentage of bleached leaf area (F

## Supplementary Materials

Figure S1: VIGS infection in various sunflower genotypes. The red asterisk identifies plants used to calculate the percentage of bleached leaf area (Figure 3); Figure S2: Electrophoresis and plant materials used for the RT-PCR analysis across different levels of VIGS-infected sunflower (Figure 4). Above, gel electrophoresis showing the results of RT-PCR from green tissues (samples marked with G) and tissues that had photo-bleaching symptoms (samples marked with B). The numbers shows the number of level from which the samples were collected. Below, the corresponding leaves from which samples were collected for the RT-PCR analysis. Arrows show the region of the leaf where the samples were collected. Under each leaf are the actual plant materials that have been used for RNA extraction; Table S1: Phenotyping data collected from the VIGS experiment of studying the effect of Agrobacterium co-cultivation time and vacuum application. This data was used to construct Table; Video S1: A time-lapse illustrating the changes in photo-bleached spots in a VIGS-infected sunflower over a 4-day period. A digital magnification is applied to both a mature leaf and a young leaf from the same plant. It is obvious that the photo-bleached spot on the young leaf is expanding in size, whereas in the mature leaf, the photo-bleached spot shows no changes in size or shape.

## Author Contributions

Conceptualization, I.K.; methodology, M.M., M.K., E.I., V.U. and A.V.; software, M.M.; formal analysis, M.M.; resources, M.K, Y.D. and A.V.; writing—original draft preparation, M.M.; writing—review and editing, I.K.; supervision, I.K.; project administration, A.S.; funding acquisition, A.S. All authors have read and agreed to the published version of the manuscript.

## Funding

This research was funded by the Russian Science Foundation (grant No 22-64-00076).

## Conflicts of Interest

The authors declare no conflict of interest.

## Notes

### Competing Interest Statement

The authors have declared no competing interest.

