## Supplementary material for "Advancing Virus-Induced Gene Silencing in Sunflower: key factors of VIGS spreading and a novel simple protocol": Figure S1: VIGS infection in various sunflower genotypes. The red asterisk identifies plants used to calculate the percentage of bleached leaf area (F: Table S1.pdf

**Table S1.** Phenotyping data collected from the VIGS experiment of studying the effect of Agrobacterium co-cultivation time and vacuum application. This raw data was used to construct Table 1.

| Treatment | Plant | Number of levels | All leaves | Symptomatic leaves | SL/AL | Position of uppermost symptomatic leaf (level) |
|---|---|---|---|---|---|---|
| vacuum + 2 hr co-cultivation | **1** | 2 | 3 | 1 | 0.33 | 1 |
| | **2** | 4 | 7 | 3 | 0.43 | 3 |
| | **3** | 5 | 9 | 2 | 0.22 | 2 |
| | **4** | 4 | 8 | 4 | 0.50 | 3 |
| | **5** | 4 | 8 | 3 | 0.38 | 2 |
| | **6** | 2 | 3 | 0 | 0.00 | |
| | **7** | 3 | 6 | 6 | 1.00 | 3* |
| | **8** | 4 | 7 | 3 | 0.43 | 3 |
| | **9** | 3 | 6 | 5 | 0.83 | 3* |
| | **10** | 4 | 7 | 5 | 0.71 | 4* |
| | **11** | 3 | 6 | 0 | 0.00 | |
| | **12** | 3 | 6 | 0 | 0.00 | |
| vacuum + 6 hr co-cultivation | **1** | 4 | 7 | 0 | 0.00 | |
| | **2** | 4 | 7 | 5 | 0.71 | 4* |
| | **3** | 3 | 6 | 5 | 0.83 | 3* |
| | **4** | 3 | 5 | 3 | 0.60 | 2 |
| | **5** | 2 | 4 | 3 | 0.75 | 2* |
| | **6** | 3 | 6 | 4 | 0.67 | 3* |
| | **7** | 4 | 8 | 5 | 0.63 | 3 |
| | **8** | 3 | 6 | 0 | 0.00 | |
| | **9** | 3 | 6 | 5 | 0.83 | 3* |
| | **10** | 4 | 8 | 5 | 0.63 | 4* |
| | **11** | 4 | 7 | 0 | 0.00 | |
| | **12** | 4 | 7 | 3 | 0.43 | 4* |
| | **13** | 4 | 7 | 5 | 0.71 | 4* |
| vacuum + 18 hr co-cultivation | **1** | 3 | 6 | 5 | 0.83 | 3* |
| | **2** | 4 | 7 | 4 | 0.57 | 2 |
| | **3** | 3 | 6 | 3 | 0.50 | 2 |
| | **4** | 3 | 6 | 0 | 0.00 | |
| | **5** | 3 | 6 | 4 | 0.67 | 3* |
| | **6** | 4 | 7 | 6 | 0.86 | 4* |
| | **7** | 4 | 7 | 5 | 0.71 | 3 |
| | **8** | 4 | 7 | 1 | 0.14 | 2 |
| | **9** | 3 | 6 | 0 | 0.00 | |
| | **10** | 3 | 5 | 0 | 0.00 | |
| | **11** | 4 | 7 | 4 | 0.57 | 3 |
| | **12** | 3 | 5 | 0 | 0.00 | |
| | **13** | 5 | 9 | 0 | 0.00 | |

| | | | | | | |
|---|---|---|---|---|---|---|
| **no vacuum + 2 hr co-cultivation** | **1** | 5 | 9 | 6 | 0.67 | 3 |
| | **2** | 3 | 6 | 0 | 0.00 | |
| | **3** | 3 | 6 | 1 | 0.17 | 1 |
| | **4** | 4 | 7 | 3 | 0.43 | 4* |
| | **5** | 4 | 8 | 5 | 0.63 | 3 |
| | **6** | 3 | 5 | 0 | 0.00 | |
| | **7** | 4 | 7 | 4 | 0.57 | 3 |
| | **8** | 4 | 7 | 0 | 0.00 | |
| | **9** | 3 | 6 | 4 | 0.67 | 3* |
| | **10** | 4 | 7 | 2 | 0.29 | 2 |
| | **11** | 3 | 6 | 0 | 0.00 | |
| | **12** | 3 | 6 | 5 | 0.83 | 3* |
| | **13** | 3 | 5 | 1 | 0.20 | 1 |
| **no vacuum + 6 hr co-cultivation** | **1** | 3 | 6 | 0 | 0.00 | |
| | **2** | 4 | 8 | 3 | 0.38 | 3 |
| | **3** | 4 | 7 | 5 | 0.71 | 3 |
| | **4** | 4 | 7 | 0 | 0.00 | |
| | **5** | 3 | 6 | 0 | 0.00 | |
| | **6** | 3 | 6 | 0 | 0.00 | |
| | **7** | 3 | 5 | 4 | 0.80 | 2 |
| | **8** | 3 | 6 | 0 | 0.00 | |
| | **9** | 3 | 6 | 0 | 0.00 | |
| | **10** | 3 | 5 | 5 | 1.00 | 3* |
| | **11** | 3 | 6 | 0 | 0.00 | |
| **no vacuum + 18 hr co-cultivation** | **1** | 2 | 4 | 0 | 0.00 | |
| | **2** | 3 | 5 | 0 | 0.00 | |
| | **3** | 3 | 6 | 0 | 0.00 | |
| | **4** | 3 | 6 | 0 | 0.00 | |
| | **5** | 3 | 5 | 0 | 0.00 | |
| | **6** | 3 | 6 | 0 | 0.00 | |
| | **7** | 3 | 6 | 0 | 0.00 | |
| | **8** | 4 | 7 | 4 | 0.57 | 3 |
| | **9** | 4 | 7 | 3 | 0.43 | 2 |
| | **10** | 3 | 5 | 0 | 0.00 | |
| | **11** | 3 | 6 | 5 | 0.83 | 2 |
| | **12** | 3 | 5 | 0 | 0.00 | |
| | **13** | 2 | 4 | 0 | 0.00 | |
| | **14** | 2 | 3 | 0 | 0.00 | |
| | **15** | 3 | 6 | 0 | 0.00 | |

* - These plants has photo-bleaching symptoms on the highest leaf (uppermost symptomatic leaf = highest leaf)
