## Supplementary figures and images for "Advancing Virus-Induced Gene Silencing in Sunflower: key factors of VIGS spreading and a novel simple protocol"

### Figure S1.jpg

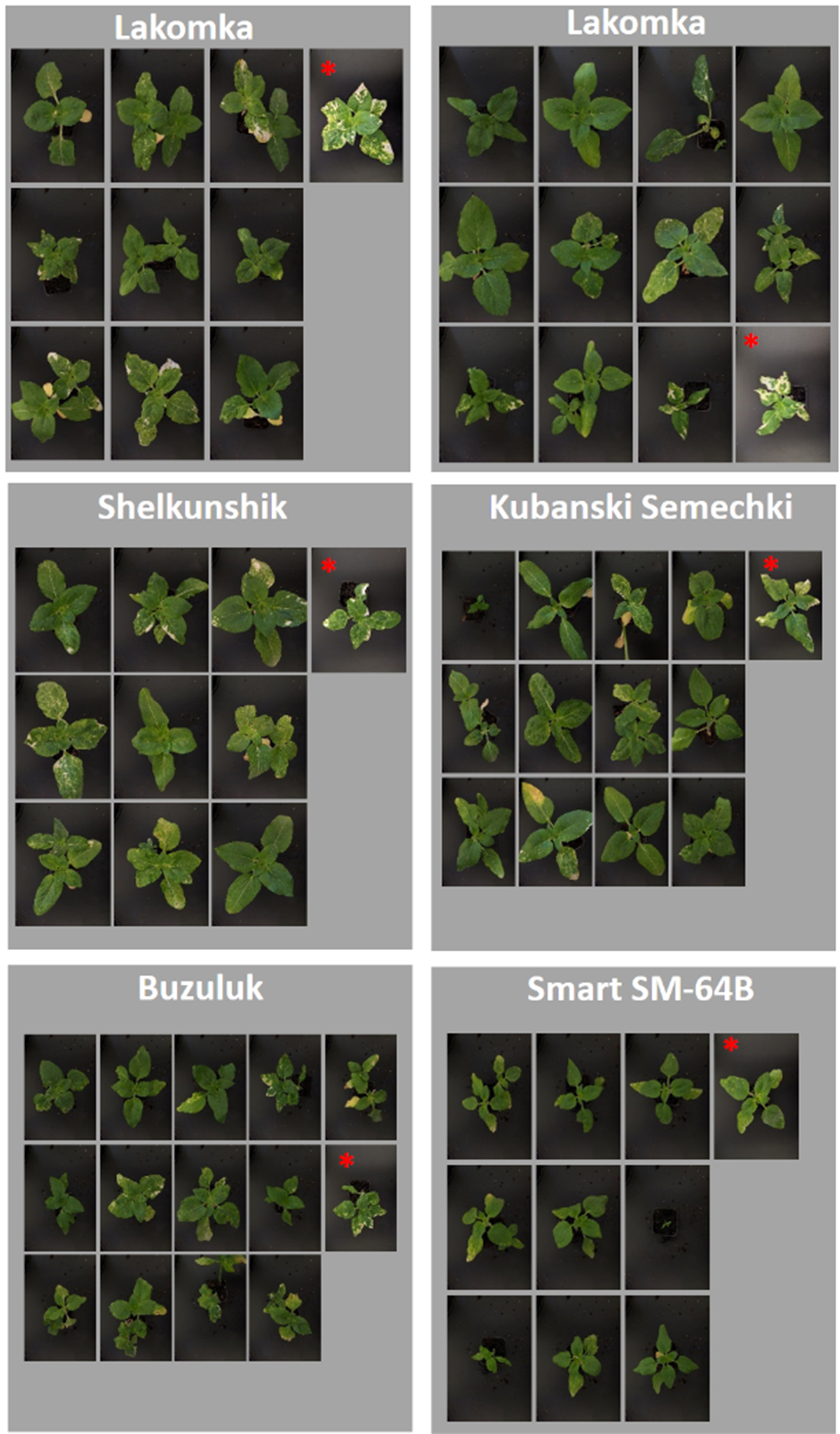

### Figure S2.jpg

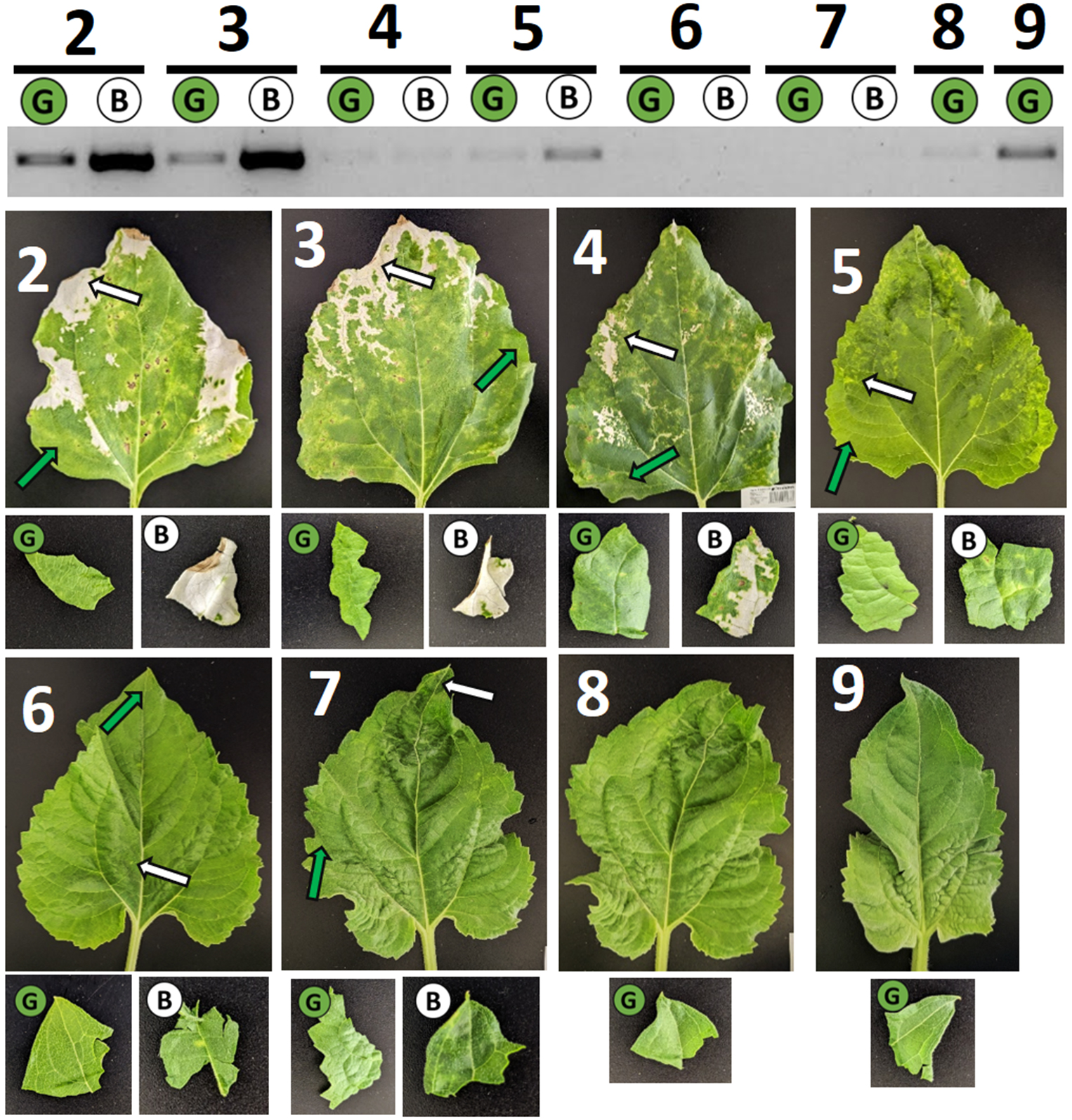
